# Partitioning the Impacts of Spatial-Temporal Variation in Demography and Dispersal on Metapopulation Growth Rates

**DOI:** 10.1101/2023.11.01.565238

**Authors:** Sebastian J. Schreiber

## Abstract

Spatial-temporal variation in environmental conditions is ubiquitous in nature. This variation simultaneously impacts survival, reproduction, and movement of individuals and, thereby, the rate at which metapopulations grow. Using the tools of stochastic demography, the metapopulation growth rate is decomposed into five components corresponding to temporal, spatial, and spatial-temporal variation in fitness, and spatial and spatial-temporal covariation in dispersal and fitness. While temporal variation in fitness always reduces the metapopulation growth rate, all other sources of variation can either increase or reduce the metapopulation growth rate. Increases occur either by reducing the impacts of temporal variation or by generating a positive fitness-density covariance where individuals tend to concentrate in higher-quality patches. For example, positive auto-correlations in spatial-temporal variability in fitness generate this positive fitness-density covariance for less dispersive populations, but decrease it for highly dispersive populations, e.g. migratory species. Negative auto-correlations in spatialtemporal variability have the opposite effects. Positive covariances between movement and future fitness, on short or long time scales, increase growth rates. These positive covariances can arise is unexpected ways. For example, the win-stay, lose-shift dispersal strategy in negatively autocorrelated environments can generate positive spatial covariances that exceed negative spatial-temporal covariances. This decomposition of the metapopulation growth rate provides a way to quantify the relative importance of fundamental sources of variation on metapopulation persistence.

## Introduction

All populations experience spatial and temporal variation in environmental conditions. Within populations, this variation generates individual variation in survival, growth, and fecundity across space and time. Faced with these demographic consequences, individuals may disperse between habitat patches and, thereby, alter their demographic fates. As environmental conditions vary across time, so may patterns of dispersal. This covariation in demography and dispersal in space and time can impact the rate at which populations grow or decline. Many studies have identified important ways that this covariation impacts population growth [Hastings, 1983, Levin et al., 1984, Wiener and Tuljapurkar, 1994, Jansen and Yoshimura, 1998, Schmidt, 2004, Roy et al., 2005, Schreiber and Lloyd-Smith, 2009, Schreiber, 2010, Evans et al., 2013, Williams and Hastings, 2013, Snyder et al., 2014, Crone, 2016]. However, as these studies typically focused on only one or two sources of variation, gaps remains in our understanding of these impacts and the relative importance of these different sources of variation for population growth rates.

Temporal and spatial variation in demography, in and of themselves, can have opposing effects on population growth. For unstructured populations (e.g. no spatial structure), temporal variation in survival or fecundity reduces population growth rates [Lewontin and Cohen, 1969, Tuljapurkar, 1990, Boyce et al., 2006]. This reduction stems from population growth being a multiplicative process and reductions in survival or fecundity having a larger impact on population growth than comparable increases in these vital rates. For sedentary populations, spatial variability in survival or fecundity always increases population growth rates [Karlin, 1976, Hastings, 1983, Schreiber and Lloyd-Smith, 2009, Snyder and Chesson, 2003, Cantrell et al., 2017]. This positive effect stems from populations growing most rapidly where fitness is highest and individuals rarely dispersing elsewhere. As the population concentrates in locations with high fitness, the covariance between population density and fitness becomes positive i.e. a positive fitness-density covariance [Chesson, 2000]. Crone [2016] illustrated these theoretical conclusions using integral projection models for the prairie forb, *Pulsatilla patens*. For these relatively sedentary populations, Crone [2016] found that “environmental stochasticity reduced population growth rates relative to the average, whereas spatial heterogeneity increased population growth rates.” For highly mobile populations, theory predicts that the positive effect of spatial variability on population growth is reduced and possibly reversed [Karlin, 1976, Hastings, 1983, Kirkland et al., 2006, Schreiber and Lloyd-Smith, 2009, Altenberg, 2012a,b]. For example, following rapid environmental change, individuals may mistakenly prefer patches with lower fitness. This generates a negative fitness-density covariance and reduces the population growth rate [Battin, 2004, Schreiber and Lloyd-Smith, 2009, Hale and Swearer, 2016]. However, the exact conditions under which spatial variation, in general, reduces population growth rates are not known.

The simultaneous impacts of spatial and temporal variation in demography on population growth are more nuanced. For spatially structured populations, serially uncorrelated temporal variation in demographic rates still reduce population growth rates [Tuljapurkar, 1990, Boyce et al., 2006]. However, if this temporal variation is spatially asynchronous, dispersal can act as a spatial bet-hedging strategy and neutralize the negative impact of temporal variation on population growth [Kuno, 1981, Metz et al., 1983, Wiener and Tuljapurkar, 1994, Jansen and Yoshimura, 1998, Evans et al., 2013, Kortessis et al., 2023, Jaggi et al., 2024]. Notably, populations coupled by dispersal can have positive growth rates despite all local growth rates being negative [Metz et al., 1983, Jansen and Yoshimura, 1998, Evans et al., 2013, Jaggi et al., 2024]. In contrast to serially uncorrelated temporal variation, positively auto-correlated fluctuations in survival or reproduction can increase growth rates of moderately dispersing populations [Roy et al., 2005, Schreiber, 2010, Kortessis et al., 2020]. Intuitively, positive auto-correlations imply that higher quality patches at one moment tend to remain higher quality. Population densities rapidly build up in these patches and dispersal seeds other patches to capitalize on shifting environmental conditions. In the long term, this results in population densities accumulating in higher quality patches and higher population growth rates [Roy et al., 2005, Kortessis et al., 2023]. Consistent with these theoretical predictions, experiments with *Paramecium aurelia* found that temporally autocorrelated fluctuations in temperature and moderate dispersal rates led to positive population growth rates despite negative growth rates on average [Matthews and Gonzalez, 2007]. These studies, however, didn’t consider the effects of spatial heterogeneity or negatively auto-correlated fluctuations on population growth.

Variation in environmental conditions can drive variation in rates of active or passive dispersal [Schmidt, 2004, Haugen et al., 2006, Ellner and Schreiber, 2012, Williams and Hastings, 2013, Snyder et al., 2014, Catalano et al., 2021, Mason et al., 2022, Peniston et al., 2023]. Active dispersers may use environmental cues to determine whether or not to leave their current patch [Schmidt, 2004, Clobert, 2012, Carroll et al., 2018]. For example, little penguins (*Eudyptula minor*) exhibit behavior consistent with the win-stay, lose-shift rule. Individuals were less likely to leave sites where they had greater foraging success and more likely to leave sites where foraging was less successful [Carroll et al., 2018]. Simulating mathematical models, Schmidt [2004] found that the win-stay, lose-shift rule combined with positively auto-correlated fluctuations in site quality increased population growth rates. Intuitively, with positive autocorrelations, site quality tends to persist over time. Hence, by remaining in higher quality sites and leaving lower quality sites, individuals tends to concentrate in higher quality sites and increase the population growth rate.

Movement of passive dispersers varies with the speed and direction of environmental currents (e.g. air, wind), their dispersal vectors, and their physiological state [O’Connor et al., 2007, Clobert, 2012, Catalano et al., 2021, Mason et al., 2022, Peniston et al., 2023]. For example, pelagic larvae are transported by spatially and temporally varying ocean currents [Catalano et al., 2021] and the time spent riding these currents is temperature dependent [O’Connor et al., 2007]. Theory predicts that this temporal variation in habitat connectivity can reduce or inflate population growth rates [Williams and Hastings, 2013, Snyder et al., 2014]. However, as these theories consider specific scenarios (i.e. two patches with no demographic variation, or serially uncorrelated fluctuations), they do not provide general principles explaining how variable dispersal impacts population growth rates.

Collectively, these earlier studies highlight the complex ways that spatial and temporal variation in demography and dispersal can impact population growth. Despite significant progress, there remain several gaps remain in our understanding of the impacts of certain forms of variation. More importantly, the relative importance of these different sources of variation on population growth is not well understood. To address these issues, I analyze a class of stochastic matrix models with any number of patches and accounting for many forms of spatial-temporal variation in demography and dispersal. The analysis provides a method to decompose the population growth rate into six contributions due to the average fitness, temporal, spatial, and spatial-temporal variation in fitness, and spatial and spatial-temporal covariation in fitness and dispersal. Using this decomposition one can evaluate the relative importance of different sources of variation for metapopulation persistence.

## Model and methods

### A Metapopulation Model

The model represents a metapopulation living in *k* patches coupled by dispersal. These patches may correspond to geographically, distinct patches of one or more habitat types, or discretizations of a continuous landscape. The population density in the *i*-th patch is *n*_*i,t*_ at time *t*. The population census times *t* = 0, 1, 2, 3, … are discrete. Between successive censuses, I assume that survival and reproduction are followed by dispersal. This ordering, however, is not critical for the main results. Survival and reproduction in patch *i* changes the local population density by a multiplicative factor *λ*_*i,t*_ at time *t*. Even though the model doesn’t assume that individuals only live for one time step, I call this multiplicative factor *λ*_*i,t*_ the fitness of an individual in patch *i* at time *t*. After survival and reproduction, a fraction *d*_*ij,t*_ of individuals from patch *j* disperse to patch *i*. Summing over all the contributions of individuals to each patch determines the metapopulation dynamics:

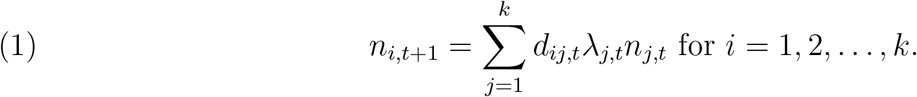

For mathematical convenience, I assume all dispersing individuals arrive in some patch i.e.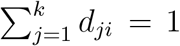. To account for mortality due to dispersal, one can reduce the fitness in each patch by the fraction dying due to dispersal i.e. if a fraction *δ*_*ij,t*_ die dispersing from patch *j* to *I* then *λ*_*j,t*_ becomes 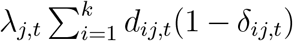 and *d*_*ij,t*_ becomes 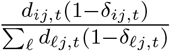. While this model does not include density-dependence, one can view it as a linearization of a density-dependent model about the extinction equilibrium. This linearization determines how metapopulations grow at low population densities and, consequently, whether the metapopulation persists or not [Benaïm and Schreiber, 2009, Roth and Schreiber, 2014, Benaïm and Schreiber, 2019].

One can rewrite the model (1) in a simpler form using vector-matrix notation. The vector of population densities at time *t* is

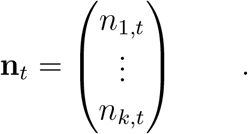

Survival and reproduction at time *t* corresponds to multiplying by the demography matrix

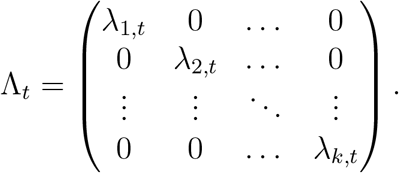

Dispersal at time *t* corresponds to multiplying by the dispersal matrix

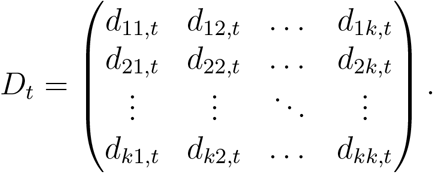

With this vector-matrix notation, the model (1) becomes

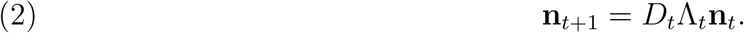

Fluctuations in environmental conditions drive the fluctuations in demography Λ_*t*_ and dispersal *D*_*t*_. The environmental state *θ*_*t*_ at time *t* lies in the set Θ of possible environmental states. This set Θ may consist of a finite number of environmental states {1, 2, …, *m*} e.g. wet and cool versus hot and dry. Alternatively, Θ may consist of continuum of states Θ ⊂ (−∞,∞)^*m*^, or some combination of continuous and discrete states. Consistent with classical stochastic demography [Ruelle, 1979, Tuljapurkar, 1990, Boyce et al., 2006], I assume that the sequence of environmental states *θ*_1_, *θ*_2_, *θ*_3_, … is (asymptotically) stationary and ergodic. For example, for a discrete environmental state space, the environmental dynamics may be given by an irreducible, aperiodic Markov chain on Θ = {1, 2, …, *m*}. Alternatively, for a continuous environmental state space, the dynamics may be given by any Markov process on a compact set that is asymptotically stationary and ergodic such as convergent multivariate auto-regressive processes, quasi-periodic motions, and certain classes of chaotic dynamics. To couple population dynamics to the environmental dynamics, the entries of the demographic and dispersal matrices are viewed as functions of the environment. Namely, in environment *θ*, the demography matrix is the diagonal matrix Λ(*θ*) with diagonal entries *λ*_*i*_(*θ*) and the dispersal matrix is **D**(*θ*) with entries *d*_*ij*_(*θ*). With this coupling to the environmental dynamics, the demographic and dispersal matrices at time *t* in (2) are

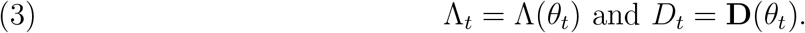

Under suitable technical assumptions, Ruelle [1979] provided that there is quantity *r* characterizing the long-term growth rate of the metapopulation. Specifically, if

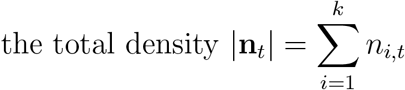

at time *t*, then the metapopulation growth rate 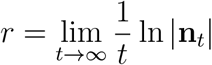 (with probability one) whenever |**n**_0_| *>* 0.

Roughly, the total metapopulation population density, **n**_*t*_, grows exponentially at rate *r*, increasing when *r >* 0 and decreasing when *r <* 0. The sign of the metapopulation growth rate *r* has implications for models accounting for density-dependence or demographic stochasticity. For models with negative density-dependence, positive values of *r* imply the metapopulation persists, while negative values imply the metapopulation approaches extinction exponentially quickly [Benaïm and Schreiber, 2009, Benaïm and Schreiber, 2019]. For density-independent models with demographic stochasticity (i.e. a multi-type branching processes in a random environment [Athreya and Ney, 2004]), *r >* 0 implies that there is a positive probability of metapopulation establishment, while *r* ≤ 0 implies establishment is impossible.

### Decomposing the sources of variation

In order to identify the relative roles of spatial and temporal variation on the metapopulation growth rate *r*, I decompose the demography and dispersal matrices into components corresponding to temporal variation, spatial variation, or spatial-temporal variation. Averaging the fitnesses 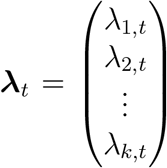 across space at time *t* yields the scalar the spatial average of fitness at time 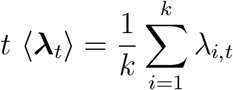.

Averaging the fitnesses across time yields the vector the temporal averages of fitnesses 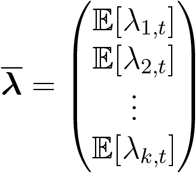 where 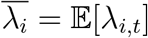 denotes the expected fitness in patch *i*. By stationarity of the environmental process, these expected fitnesses correspond to the long-term temporally averaged fitnesses and, therefore, do not depend on time *t* i.e. 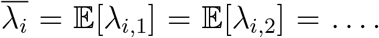 Averaging simultaneously across space and time yields the spatial-temporal average of fitness 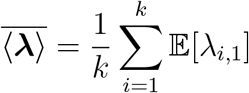.

Using these averages, the fitnesses ***λ***_*t*_ can be decomposed into four components. The first component, 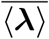, is the average fitness experienced by the population across space and time. Temporal, spatial, and spatial-variability in the fitnesses are quantified as deviations from this spatial-temporal average. Subtracting the spatial-temporal average from spatial average in year *t*, 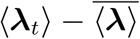, yields the temporal deviation in year *t*. Positive temporal deviations correspond to years where individuals, on average across space, have higher fitness. Subtracting the spatialtemporal average from the vector of temporal averages, 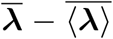, yields the vector of spatial deviations. Positive components of this vector correspond to patches that, on average across time, have higher fitness. The remaining deviations in fitness are the vector of spatial-temporal deviations

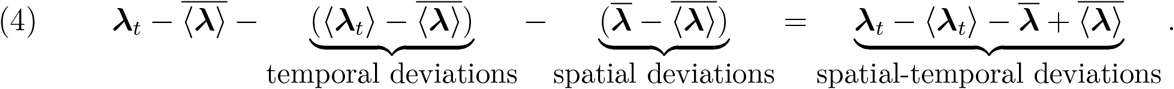

Positive components of this vector correspond to years of higher fitness for a given patch. Rearranging (4) yields the following decomposition of the demography:

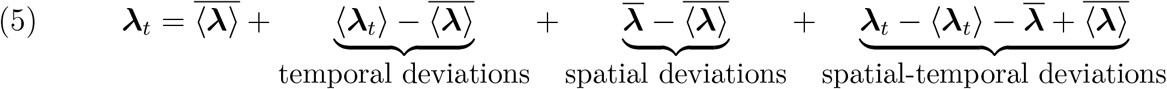

This decomposition of the demography is similar to decompositions used by Johnson and Hastings [2023] and Kortessis et al. [2023].

To quantify variation in dispersal, I assume there is a symmetric dispersal matrix *D*_sym_ (i.e. *d*_*ij*_ = *d*_*ji*_ for all *i* ≠ *j*) around which the spatial variation in dispersal occurs. For two patch models, this symmetric matrix is

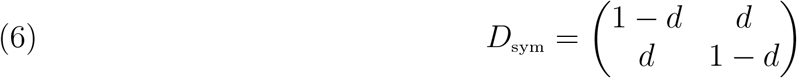

where *d* is the probability an individual disperses to the other patch. For models with more patches, this symmetric matrix could represent a random walk or diffusion with any pattern of connectivity between the patches [Chung, 1997, Spielman, 2019]. There are two advantages of using a symmetric matrix *D*_sym_ as a frame of reference. First, in the absence of spatial variation in the fitnesses, symmetric dispersal ensures that, in the long term, individuals are uniformly distributed across the patches i.e. there is no spatial heterogeneity in population densities. Second, symmetric matrices have an orthogonal basis of eigenvectors providing a natural way to decompose spatial variation using a generalization of discrete Fourier analysis [Stein and Shakarchi, 2011]. While an approximation of the metapopulation growth rate *r* is possible when only the first condition holds (i.e. *D*_sym_ is doubly stochastic but not symmetric), the biological interpretation is more complex (see, for example, equation (B4) in Appendix B).

Variation of the dispersal matrix *D*_*t*_ around the reference dispersal matrix *D*_sym_ takes two forms: spatial variation and spatial-temporal variation. The spatial variation corresponds to the difference between the time-averaged dispersal matrix 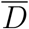 and the reference matrix *D*_sym_. A positive entry in this difference corresponds to increased movement, on average over time, between a pair of patches. Unlike demography, there is no natural way to define pure temporal variation in *D*_*t*_. This stems from the spatial average of the entries of *D*_*t*_ not varying over time i.e. 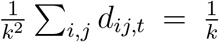 for all *t*. Intuitively, movement into one patch is balanced by a corresponding movement out of another patch. Hence, beyond the spatial deviations, there are only the spatial-temporal deviations yielding the decomposition

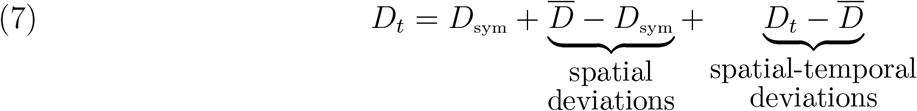

Positive entries in the spatial-temporal deviations 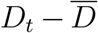 correspond to years with more movement from one patch to another than in a typical year.

### Analytical and numerical approximations of the metapopulation growth rate *r*

Finding explicit analytic expressions for the metapopulation growth rate *r* is generally intractable. However, using methods of stochastic demography [Ruelle, 1979, Tuljapurkar, 1990], I derive an analytical approximation of *r* when the temporal and spatial variations in demography and dispersal are small. To validate the accuracy of these analytical approximations, I numerically compute *r* using the power method [Caswell, 2001]. Schreiber [2024] provides R code for doing the analytical and numerical approximations, and recreating the figures.

To derive the small variance approximation in a mathematically rigorous manner, I use the work of Ruelle [1979] who proved that *r* is an analytic function of the environment-demography mapping *θ* → **D**(*θ*)Λ(*θ*) =: **A**(*θ*). Ruelle [1979]’s proof of this fact provides an approach to explicitly compute all derivatives of *r* as a function of the environment-demography mapping. In particular, this approach provides an alternative, more direct method to compute the first order and second order approximations of Tuljapurkar [1990]. Furthermore, Ruelle [1979]’s approach also allows one to derive second-order mixed derivatives of *r* that are used to quantify the interactive effect of covariation in dispersal and demography on *r*. Using Ruelle [1979]’a approach, Appendix A derives these first-order and second-order derivatives for general stochastic matrix models. Using these derivative formulas, Appendix B derives a second order Taylor expansion for *r* around the unperturbed model 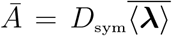. The Taylor expansion consists of sum of three terms: the zero-th order term ln 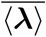, a second-order correction term due to demographic variation, and a second-order mixed correction term due co-variation between demography and dispersal.

To approximate *r* numerically, I use the power method [Caswell, 2001] where one re-normalizes the matrix dynamics in (2) at each time step:

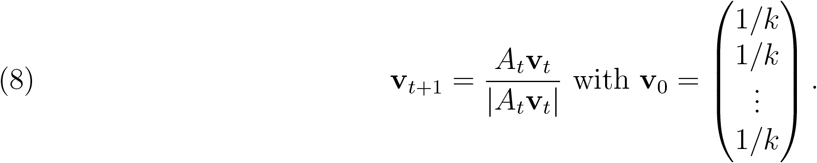

The work of Ruelle [1979] implies that

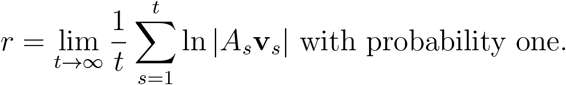

Hence,

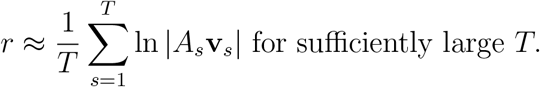

I use *T* = 100, 000 time steps for these numerical approximations.

#### The role of the fitness-density covariance

The work of Ruelle [1979] implies that **v**_*t*_ converges to a stationary sequence **v**_*t*_ representing the spatial-temporal distribution of individuals in the population in the long term. Notably, the metapopulation growth rate equals

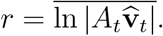

This equation provides a link to the fitness-density covariance of Chesson [2000]. Specifically, if

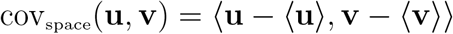

denotes the spatial covariance of two arbitrary vectors **u, v**, then

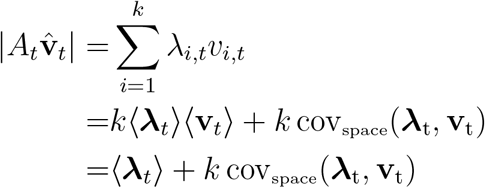

where the first line follows from the dispersal matrices being column stochastic, the second line follows from the definition of the spatial covariance, and the final line follows from (**v**_*t*_) = 1*/k* i.e. **v**_*t*_ = 1. The expression *k* cov(***λ***_t_, **v**_t_) corresponds to the spatial covariance between the fitnesses at time *t* and the (normalized) population densities at time *t*. Chesson [2000] calls this expression, the fitness-density covariance. Putting these pieces together, one arrives at the following representation of the metapopulation growth rate *r*

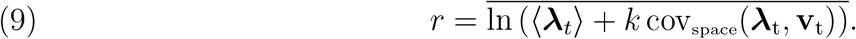

Roy et al. [2005] derived a similar formula to (9).

## Results

The main result is an analytical formula for how spatial and temporal variation in demography and dispersal contribute to the metapopulation growth rate *r*. This formula consists of two main components: the contribution of demographic variation and an interactive effect demographic and dispersal variation. To highlight the key features of these two components, I present them first for the case of *k* = 2 patches and then extend to the case of any number of patches. I conclude by deriving a relationship between the fitness-density covariance of Chesson [2000] and the metapopulation growth rate decomposition.

### Contributions of demographic variation to *r* for two patches

In case of *k* = 2 patches without dispersal variation, the dispersal matrix *D*_*t*_ = *D*_sym_ is characterized by the probability *d* that an individual disperses to the other patch, see equation (6). In the absence of births and deaths, individuals dispersing according to this dispersal matrix will ultimately by equally distributed between the two patches i.e. the stable spatial distribution is 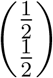. Deviations away from this spatially homogeneous distribution decay exponentially like (1 −2*d*)^*t*^ as *t*→ ∞. This decay occurs more quickly when the fraction dispersing is closer to 50%. Moreover, this decay occurs monotonically when no more than 50% of the population is dispersing (*d <* 0.5). In this case, I call the population *philopatric* (home-loving). When the fraction dispersing is greater than 50% (*d >* 0.5), I call the population *miseopatric* (home-hating). The spatial distribution of miseopatric populations exhibits dampened oscillations when approaching a spatially homogeneous state; the patch with the greatest number of individuals alternating between the two patches from time step to time step.

In the presence of spatially and temporally varying fitnesses, the metapopulation experiences a temporal variance 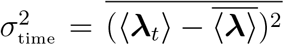 in its spatially averaged fitnesses with coefficient of variation 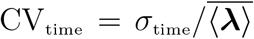, and a spatial variance 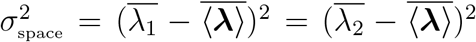 in its temporally averaged fitnesses with coefficient of variation 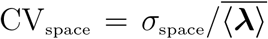. In the case of *k* = 2 patches, the entries of the vector of spatial-temporal deviations (final term in equation (5)) always have opposite signs; the spatial-temporal deviations are

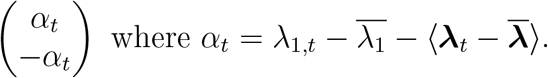

*α*_*t*_ measures the direction and magnitude of the spatial asynchrony between the two patches. When *α*_*t*_ is zero, fitnesses in both patches are equally higher or equally lower than their respective temporal averages. When *α*_*t*_ is positive, fitness in patch 1 is higher than its average fitness while fitness in patch 2 is correspondingly lower than its average fitness. Let 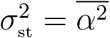 be the temporal variation in the spatial asynchrony and 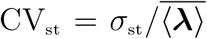 the associated coefficient of variation. The ratio of variances, 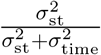, is a metric of spatial asynchrony: zero corresponds to perfect spatial synchrony while one corresponds to perfect spatial asynchrony. Let 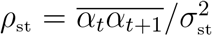 be the temporal autocorrelation in the *α*_*t*_. When the autocorrelation *ρ*_st_ is positive, time steps in which the fitness in patch 1 is greater than its average tend to be followed by time steps where the fitness in patch 1 continues to be higher than its average. When the autocorrelation *ρ*_st_ is negative, time steps in which the fitness in patch 1 is higher than its average tend to be followed by time steps where the fitness in patch 2 is higher than its average.

The analytical approximation of the metapopulation growth rate *r* depends on the average fitness and the temporal, spatial, and spatial-temporal deviations from this average i.e. each of the terms of the decomposition of fitness in equation (5). Specifically,

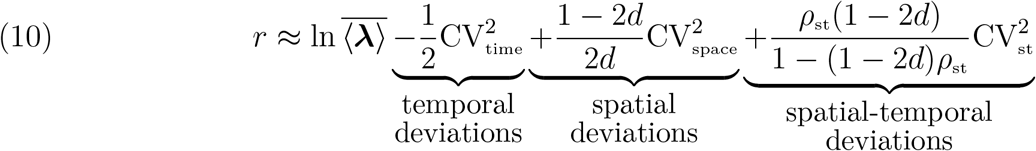

The first term ln 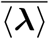 corresponds to the metapopulation growth rate in the absence of spatial and temporal variation. Consistent with classical stochastic demography, the second term, 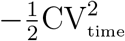, implies that temporal variation in demography reduces the metapopulation growth rate. In contrast, spatial variation (the third term) increases the metapopulation growth rate for philopatric populations (*d <* 0.5) and decreases the metapopulation growth rate for miseopatric populations (*d >* 0.5). The impact of spatially asynchronous fluctuations on the metapopulation growth rate (the fourth term) depends simultaneously on the temporal auto-correlation *ρ*_st_ and whether the populations are philopatric or miseopatric. For philopatric populations, positive autocorrelations increase the metapopulation growth rate, while negative autocorrelations reduce the metapopulation growth rate. For miseopatric populations, these impacts are reversed.

To illustrate these theoretical implications, I first consider numerical scenarios inspired by the experiments of Matthews and Gonzalez [2007] with metapopulations of *Paramecium aurelia*. By varying temperature between two values, Matthews and Gonzalez [2007] varied fitness within flasks between low (*λ*_*i,t*_ *<* 1) and high (*λ*_*i,t*_ *>* 1) values. This temperature variation was chosen so that the average fitness was less than one 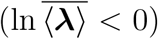. Each day (a time step), 10% of the individuals were moved between pairs of flasks (*d* = 0.1). Like their experiments, my numerical experiments varied temporal and spatial correlations in fitness (Fig. 1). In these scenarios, the analytical approximation of the metapopulation growth rate *r* closely matched the numerically estimated values of *r* (solid curves versus crosses in Fig 1A). For spatially synchronized metapopulations 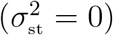, temporal variation in fitness reduced the metapopulation growth rate (yellow versus gray curves in Fig. 1A). When the patches are completely spatially asynchronous (i.e. when one patch experiences high fitness, the other experiences low fitness), the metapopulation only experiences spatial-temporal fluctuations in fitness (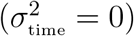); there is no reduction due to temporal variation. As the metapopulation is philopatric (*d* = 0.1), positive autocorrelations in this spatial-temporal variation increase the metapopulation growth rate, while negative autocorrelations decrease the metapopulation growth rate (red versus gray curves in Fig. 1A). Sufficiently positive autocorrelations lead to a positive metapopulation growth rate despite both patches being sinks (red curve above dashed line in Fig. 1A). When the patches are spatially independent, the joint effects of temporal variation and spatial-temporal variation result in intermediate metapopulation growth rates (blue versus yellow and red curves in Fig. 1A). This stems from spatial independence simultaneously reducing the temporal variation and the spatial-temporal variation terms of the metapopulation growth rate decomposition (10). To illustrate these implications for populations experiencing density-dependence, I ran simulations replacing the density-independent fitnesses *λ*_*i,t*_ with density-dependent fitnesses 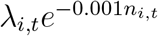. Consistent with mathematical theory [Benaïm and Schreiber, 2019], the metapopulation growth rate *r* for the density-independent model predicts the long term fate of metapopulations experiencing negative density-dependence (Fig. 1B–D), the metapopulation persisting when *r >* 0 (Fig. 1B,C) and going extinct when *r <* 0 (Fig. 1D).

**Figure 1.**
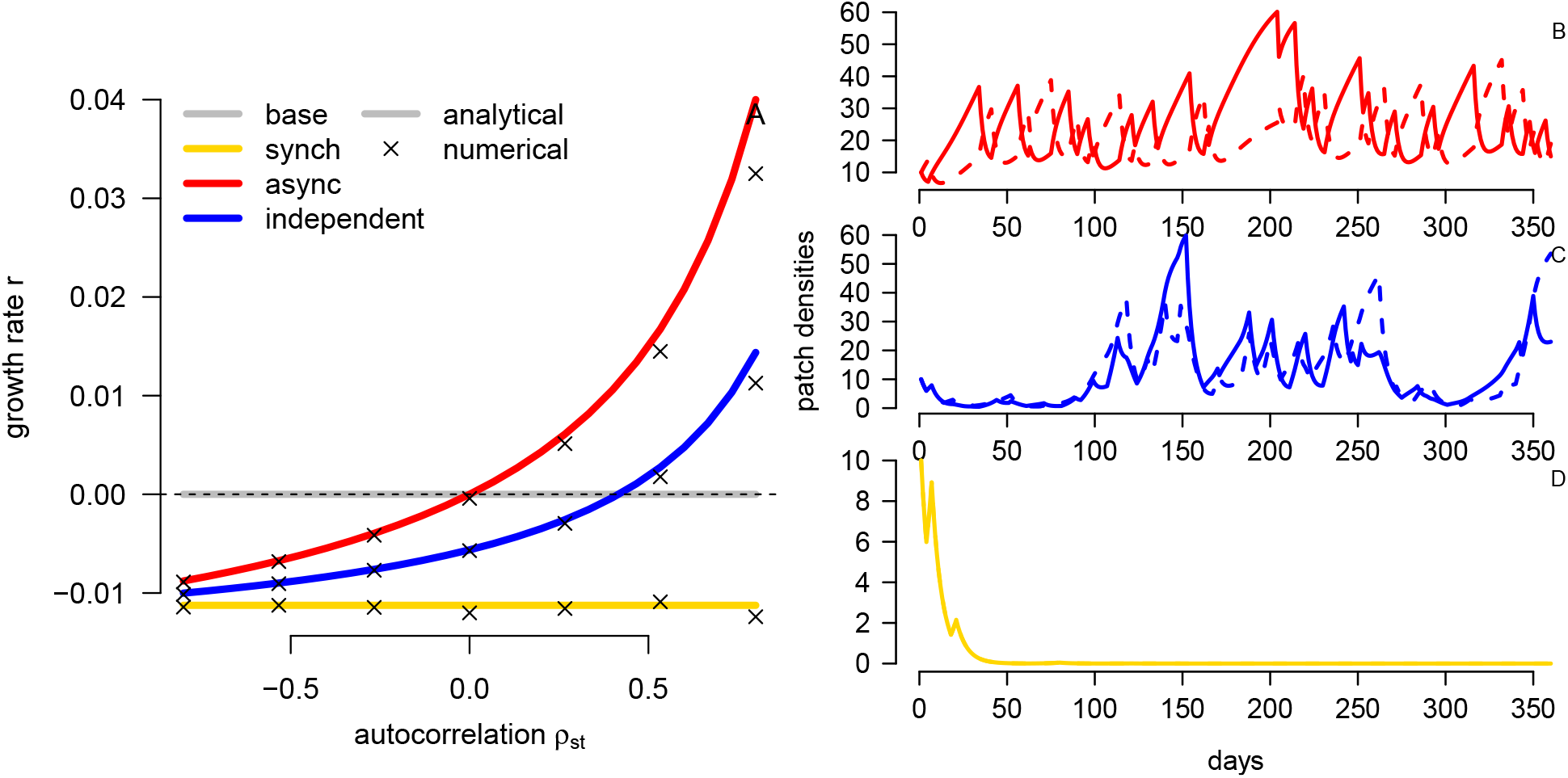
Impacts of temporal and spatial-temporal variation in fitness on two patch metapopulations. In (A), the metapopulation growth rate *r* as a function of the auto-correlation in the spatial-temporal variation. Solid curves correspond to the analytical approximation (13) and crosses correspond to numerical approximations. Spatially synchronized fluctuations in yellow, spatially asynchronous fluctuations in red, and spatially independent fluctuations in blue. The gray line corresponds to 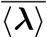. In (B)-(D), population dynamics with negative density-dependence added when *ρ*_st_ = 0.8. Different colors as described for (A). Parameters: *λ*_*i,t*_ = 0.85 and 1.15 with equal likelihood, and *d* = 0.1. For the spatially synchronous (respectively, asynchronous) models there are two environmental states corresponding to *λ*_*i,t*_ = 1.15 in both patches or *λ*_*i,t*_ = 0.85 in both patches (respectively, 1.15 in one patch, and 0.85 in the other patch). For the spatially independent model, there are four environmental states corresponding to *λ*_*i,t*_ = 1.15 in both patches, *λ*_*i,t*_ = 0.85 in both patches, and *λ*_*i,t*_ = 1.15 in one patch, *λ*_*i,t*_ = 0.85 in the other. For B-D, density-dependence is added by multiplying *λ*_*i,t*_ by exp(*−*0.001*n*_*i,t*_).

To illustrate the joint effects of spatial variation and dispersal rates on the metapopulation growth rate, I consider a scenario where patch 1 has, on average, higher but more variable fitness, while patch 2 experiences no temporal variation (Fig. 2A). While dispersal rates and temporal autocorrelations have no effect on the temporal variation component of the metapopulation growth rate *r* (not shown), the spatial and spatial-temporal components of *r* depend on both factors (Fig. 2B). For all levels of temporal autocorrelation, the spatial component of *r* decreases with the dispersal rate *d* and becomes negative when the population is miseopatric (gray dashed curve in Fig. 2B). In contrast, the effect of dispersal on the spatial-temporal component of *r* depends on the sign of the temporal autocorrelation: a positive effect for negative temporal autocorrelation (blue curve in Fig. 2B) and a negative effect for positive temporal autocorrelation (red curve in Fig. 2B). For negative temporal autocorrelations, the opposing effects of dispersal on two components of *r* result in a non-monotonic effect of dispersal on *r* (blue curve in Fig. 2A). Indeed, *r* is locally maximized at low and high dispersal rates and is minimized at intermediate dispersal rates.

**Figure 2.**
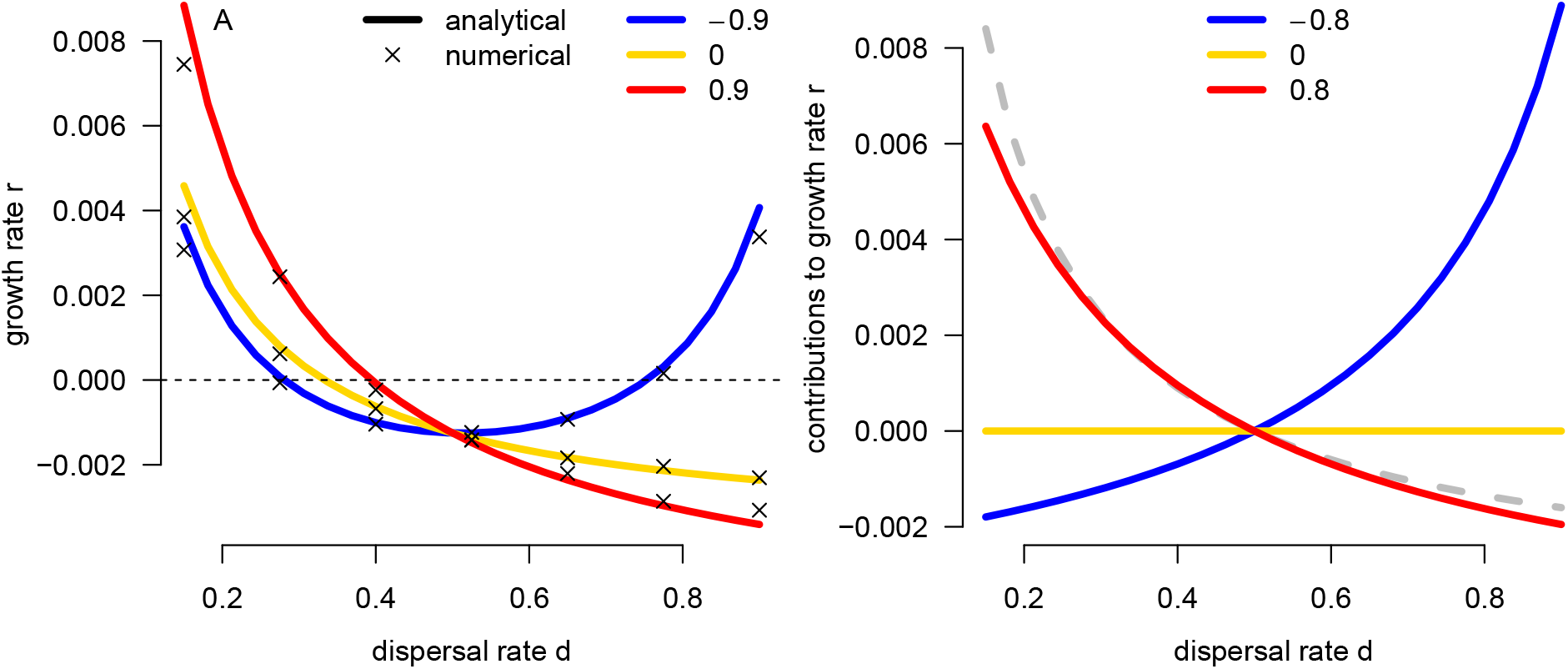
Impacts of spatial variation, temporal auto-correlation, and dispersal on the metapopulation growth rate *r*. Fitness is variable and, on average, higher in patch 1 and constant, but lower in patch 2. In (A), *r* as function of the dispersal rate *d*. Solid curves correspond to the analytical approximation (10) and crosses correspond to the numerical approximations. Different colors correspond to different values of spatial-temporal auto-correlation. In (B), the contributions of spatial variation (dashed gray) and spatial-temporal variation (yellow,red,blue) on *r*. Parameters: *λ*_*i,t*_ in patch *i* = 1 are 0.95, 1.05 with equal probability, and always 0.95 in patch *i* = 2.

### Interactive effects of dispersal and fitness variation for two patches

In the case of *k* = 2 patches, spatial and temporal variation in dispersal is determined by the changes *δ*_*i,t*_ in the fraction *d* dispersing from patch *i* = 1, 2. Namely,

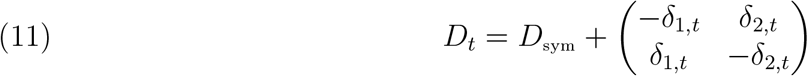

The spatial component in the dispersal matrix decomposition (7) is determined by the temporal averages 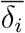 and spatial-temporal component is determined by the deviations 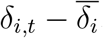.

While dispersal variation, in and of itself, has no impact on the metapopulation growth rate, it does have an impact when it covaries with demographic variation. The spatial component of this impact stems from the spatial covariance between dispersal and demography,

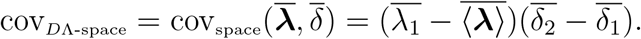

Positive values of this covariance occur when individuals, on average across time, exhibit biased movement to the patch with the higher average fitness. The impact of spatial-temporal variation on the metapopulation growth rate depends on the temporal covariance between the difference in dispersal rates between the two patches at one time step and the corresponding difference in fitness between the two patches in the next time step:

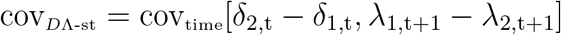

where 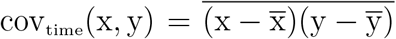 is the covariance for two stationary time series *x*_*t*_, *y*_*t*_. Let *ρ*_*D*Λ-st_ be the correlation associated with the covariance cov_*D*Λ-st_. Positive correlations occur when individuals are more likely to disperse to the patch with the higher fitness in the next time step. The effects of these spatial and spatial-temporal covariances on the metapopulation growth rate *r* are

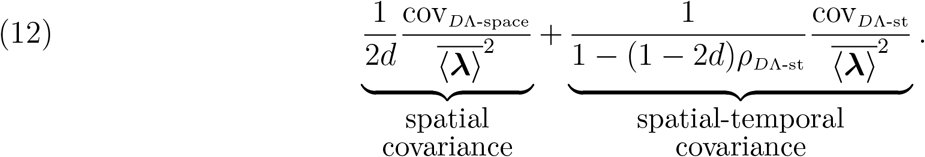

Each contribution is positive only if the corresponding covariance is positive i.e. individuals tend to move to patches of higher quality. The magnitude of these contributions decreases with the base dispersal rate *d*.

To illustrate some of the implications of the covariation of dispersal and demography on the metapopulation growth rate, Figure 3 considers a population whose individuals make use of the win-stay, lose-shift (WSLS) dispersal strategy [Schmidt, 2004]. Individuals using this strategy remain in patches that supported the highest fitness, else disperse to the other patch. Dispersing individuals (a fraction *d* of the population) utilize the WSLS strategy with probability *p*. In the numerical experiments, fitness in the patches fluctuates between lower and higher values. Patch 1, on average, supports the higher fitness, but has the lowest fitness one-quarter of the time. Greater use of the WSLS strategy (higher *p*) always increases the metapopulation growth rate (Fig 3A), but this increase is weakest when there are negative temporal autocorrelations (blue curve in Fig. 3A). At first, it may seem surprising that the WSLS strategy enhances metapopulation growth in a strongly negatively autocorrelated environment. Indeed, individuals using this strategy are more likely to end up in a patch with lower fitness in the next time step leading to a negative spatial-temporal covariance between dispersal and fitness (blue curve in Fig. 3B). However, the spatial component always has a positive effect on the metapopulation growth rate (dashed gray curve in Fig. 3B). This positive effect stems from the WSLS strategy ensuring individuals spend three-quarters of their time, on average, in the patch with the higher average fitness.

**Figure 3.**
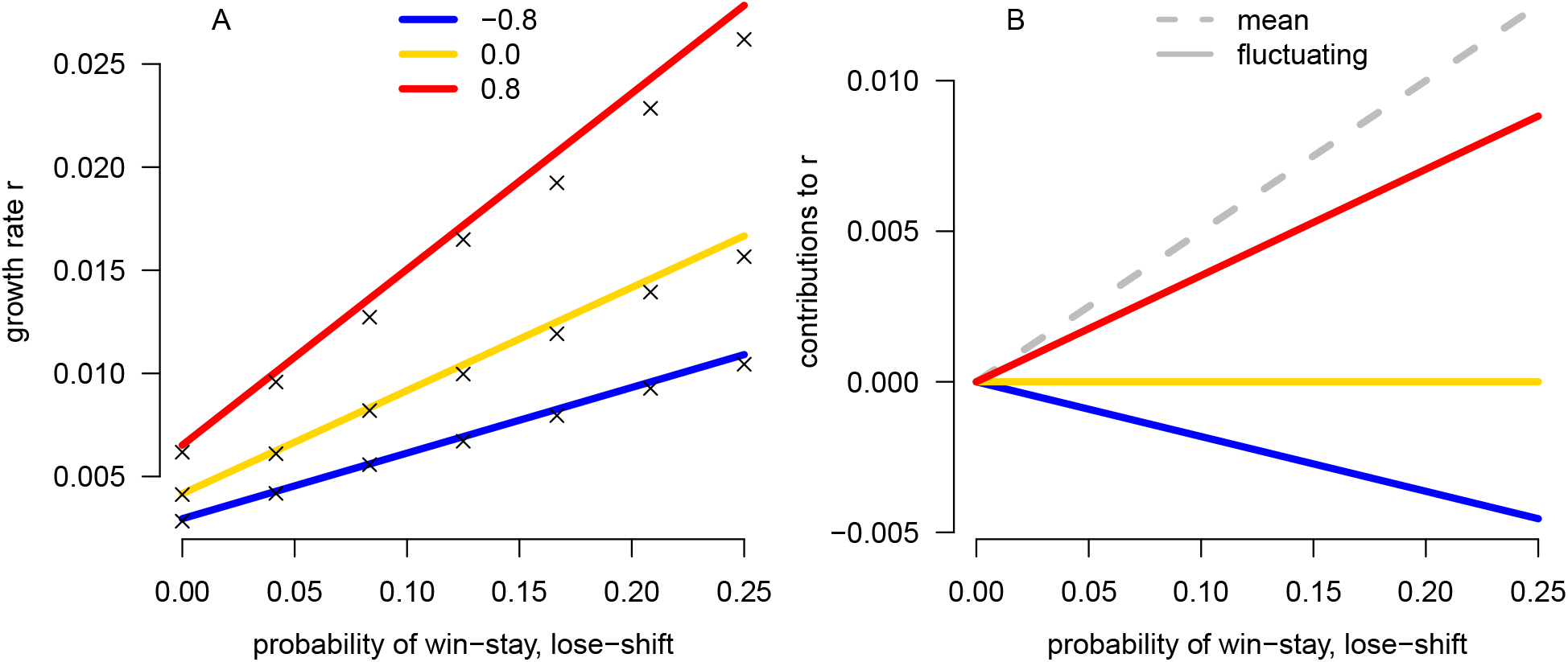
Impacts of fluctuating dispersal and demography on population growth. The fitness in patch 1 is, on average in time, higher than in patch 2, but is lower one-quarter of the time. In each time step, individuals disperse with probability *d* from patches with lower fitness and with probability *pd* from patches with higher fitness. In (A), the population growth rate plotted as function of the probability *p* of a dispersing individual playing the win-stay, lose-shift strategy. Solid curves correspond to the analytical approximation (13) and crosses correspond numerical approximations. Different colors correspond to different values of temporal auto-correlation. In (B), the contribution of the mean (dashed gray) and fluctuating (solid curves) biases in dispersal on the population growth rate. Parameters: *λ*_*i,t*_ in patch *i* = 1 are 1.2, 1.0 with equal probability, and 1.0, 0.8 in patch *i* = 2 with equal probability. Fitnesses are spatially independent and *d* = 0.3.

### Decomposition for any number of patches

In the case of *k* = 2 patches, there is only one degree of freedom for spatial variation around the temporal averages of fitness, ***λ***, corresponding to multiples of the vector 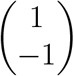. With *k* ≥ 3 patches, there are *k* −1 ≥ 2 degrees of freedom. A mathematically convenient way to characterize these *k −*1 degrees of freedom is to use a form of discrete Fourier analysis that decomposes spatial deviations from a spatially homogeneous state into a weighted sum of *k −* 1 spatial modes of variation (SMV) i.e. an orthogonal basis of subdominant eigenvectors **u**(1), **u**(2), …, **u**(*k −* 1), of the dispersal matrix *D*_sym_ normalized such that 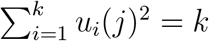. In the absence of variation in fitness or dispersal, spatial variation in the population densities along SMV **u**(*i*) decays at a fixed rate *μ*_*i*_ due to dispersal according to *D*_sym_ where *μ*_*i*_ is the eigenvalue associated with **u**(*i*). While the magnitude of *μ*_*i*_ is always less than one, its sign can be positive or negative. I call spatial modes of variation for which *μ*_*i*_ *>* 0 *philopatric spatial modes of variation*; when the initial spatial distribution of the population has all of its variation in a philopatric spatial mode, patches with higher densities will continue to have higher densities in future time steps. When the initial spatial distribution has all of its spatial variation in a SMV with negative values of *μ*_*i*_, patches with the highest densities will have the lower densities in the next time step. These are *miseopatric spatial modes of variation*. By the Gershgorin circle theorem [Horn and Johnson, 1990], miseopatric SMV only occur if there is at least one patch where at least 50% of the individuals leave i.e. there is a diagonal entry of *D*_sym_ whose value is greater than 0.5. For example, In the case of *k* = 2 patches, there is only one SMV 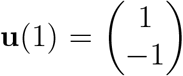 corresponding to complete spatial asynchrony. The eigenvalue associated with **u**(1) is *μ*_1_ = 1− 2*d* where *d* is the fraction of dispersing individuals.

As in the case of *k* = 2 patches, each source of variation in fitness (see (5)) and dispersal (see (7)) has a distinctive impact on the metapopulation growth rate *r*. To identify these impacts, I decompose the spatial sources of variation along the SMVs **u**(1), …, **u**(*k*). For the spatial variation in fitness, this decomposition takes the form

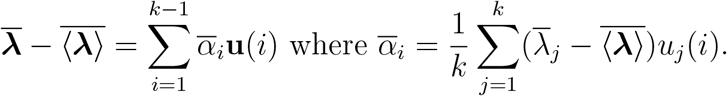

For the spatial-temporal variation, the decomposition is

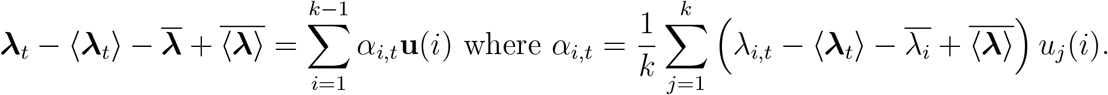

For dispersal variation, only net changes in movement for each patch impact the metapopulation growth rate

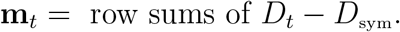

As the column sum of **m**_*t*_ always equals zero, one can express this vector as a unique linear combination of the spatial modes of variation:

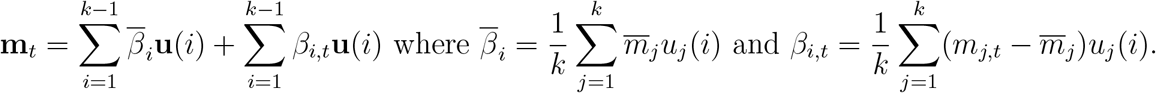

As in the two patch case, temporal variation 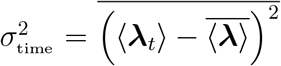 in demography reduces the metapopulation growth rate. The effect of variation along a spatial mode depends on its variance 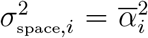 and the eigenvalue *μ*_*i*_ associated with this mode. In particular, variation along a spatial mode enhances the metapopulation growth rate only when movement along this spatial mode is philopatric i.e. *μ*_*i*_ *>* 0. The effect of spatial-temporal variation in demography depends on its variation 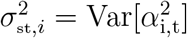 along a spatial mode and the associated temporal auto-correlation *ρ*_st_,*i* = Corr[*α*_i,t_, *α*_i,t+1_]. For philopatric spatial modes of variation (*μ*_*i*_ *>* 0), positive autocorrelations enhance metapopulation growth rates, while negative autocorrelations reduce *r*. For miseopatric spatial modes of variation, the direction of these impacts is reversed. The impact of spatial covariation in demography and dispersal on *r* depends on the dispersal-demography covariance cov_*D*Λ-space,*i*_; positive covariances along a spatial mode increase *r*. Similarly, positive spatial-temporal covariances cov_*D*Λ-st,*i*_ or auto-correlations *ρ*_*D*Λ-st,_i also increase the metapopulation growth rate. Putting all of these contributions together yields the following decomposition of the metapopulation growth rate:

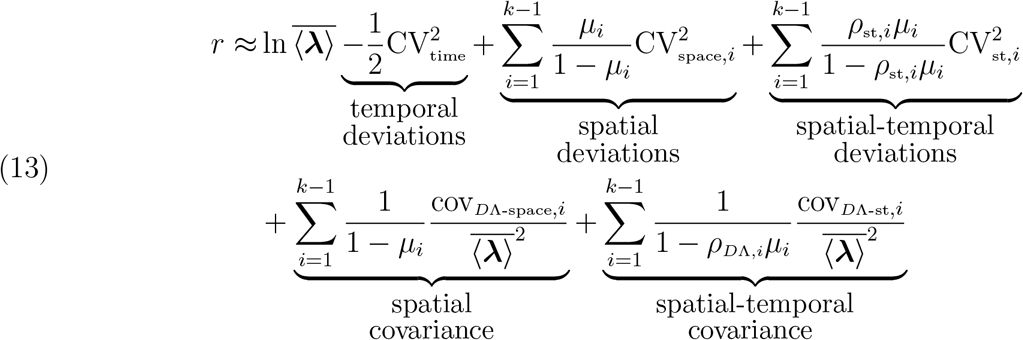

To illustrate the role of multiple spatial modes on population growth, Figures 4 and 5 consider a one-dimensional gradient of *k* = 15 patches with periodic boundary conditions. Individuals disperse preferentially to patches with higher fitness. The reference dispersal matrix *D*_sym_ corresponds to dispersal in the absence of preferences; half of the individuals in a patch move to the left, and the other half to the right. The SMVs of *D*_sym_ correspond to different spatial frequencies

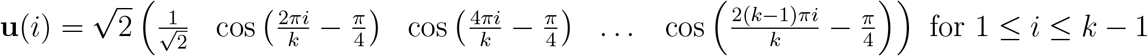

with the associated eigenvalue *μ*_*i*_ equal to 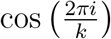 [Berlin and Kac, 1952, Demidenko, 2017]. Importantly, movement along modes corresponding to higher frequencies (i.e.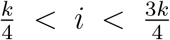) is miseopatric (i.e. *μ*_*i*_ *<* 0) and otherwise philopatric. It follows that lower versus higher frequencies of spatial variation have opposing impacts on population growth rate (Fig. 4 versus Fig. 5).

**Figure 4.**
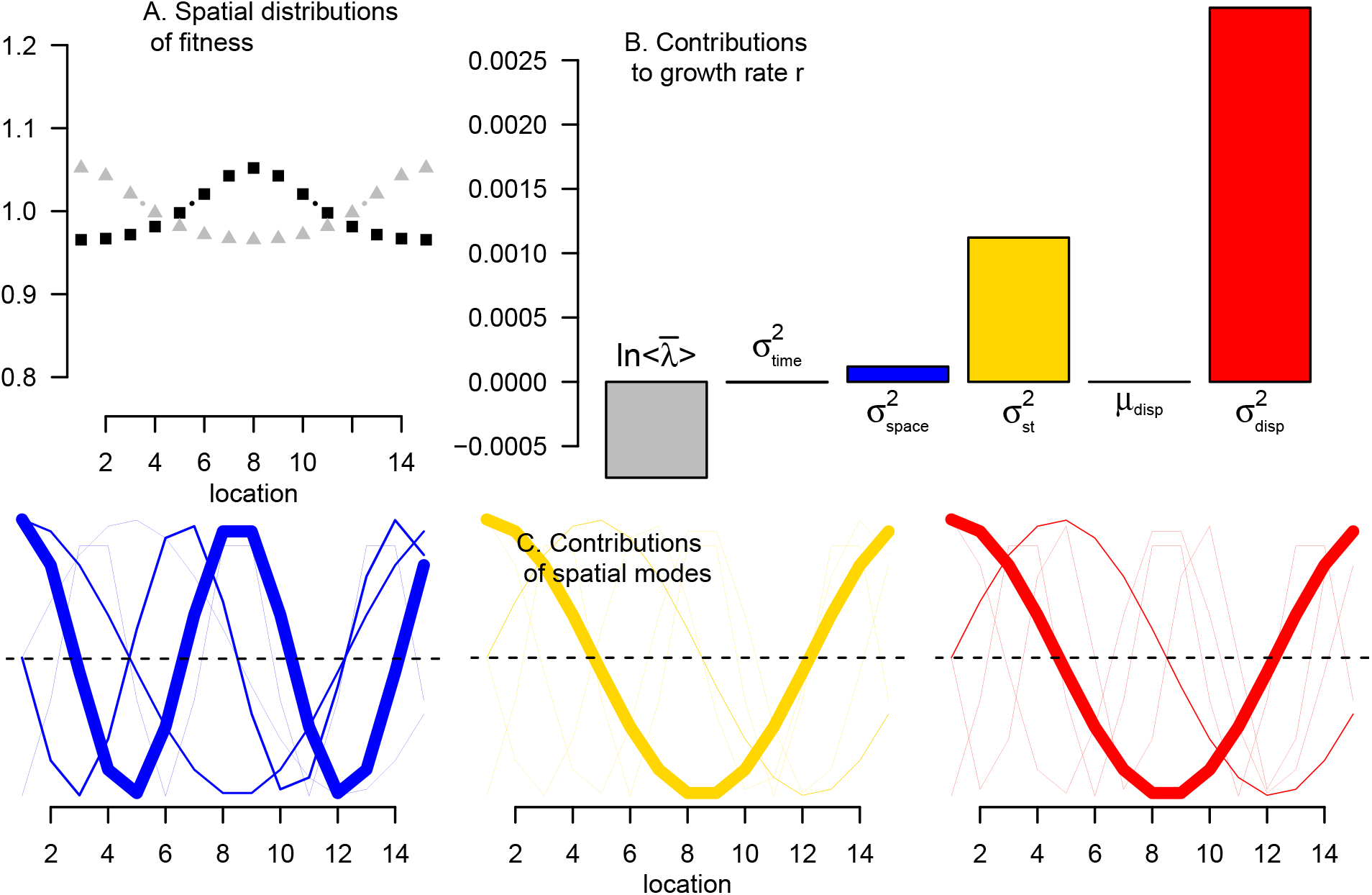
The impact of low frequency spatial variation in fitness on the metapopulation growth rate. A population living along a spatial gradient of *k* = 15 patches that varies over time between higher fitness in the center (black curve in A) and higher fitness at the edges (gray curve in A). Individuals disperse to neighboring patches with a preference for patches supporting a higher fitness. Analytical decomposition of the population growth rate in B. In C, all spatial modes of variation (SMV) plotted with the thickness of the curves proportional to the magnitude of their contribution to the metapopulation growth rate.

**Figure 5.**
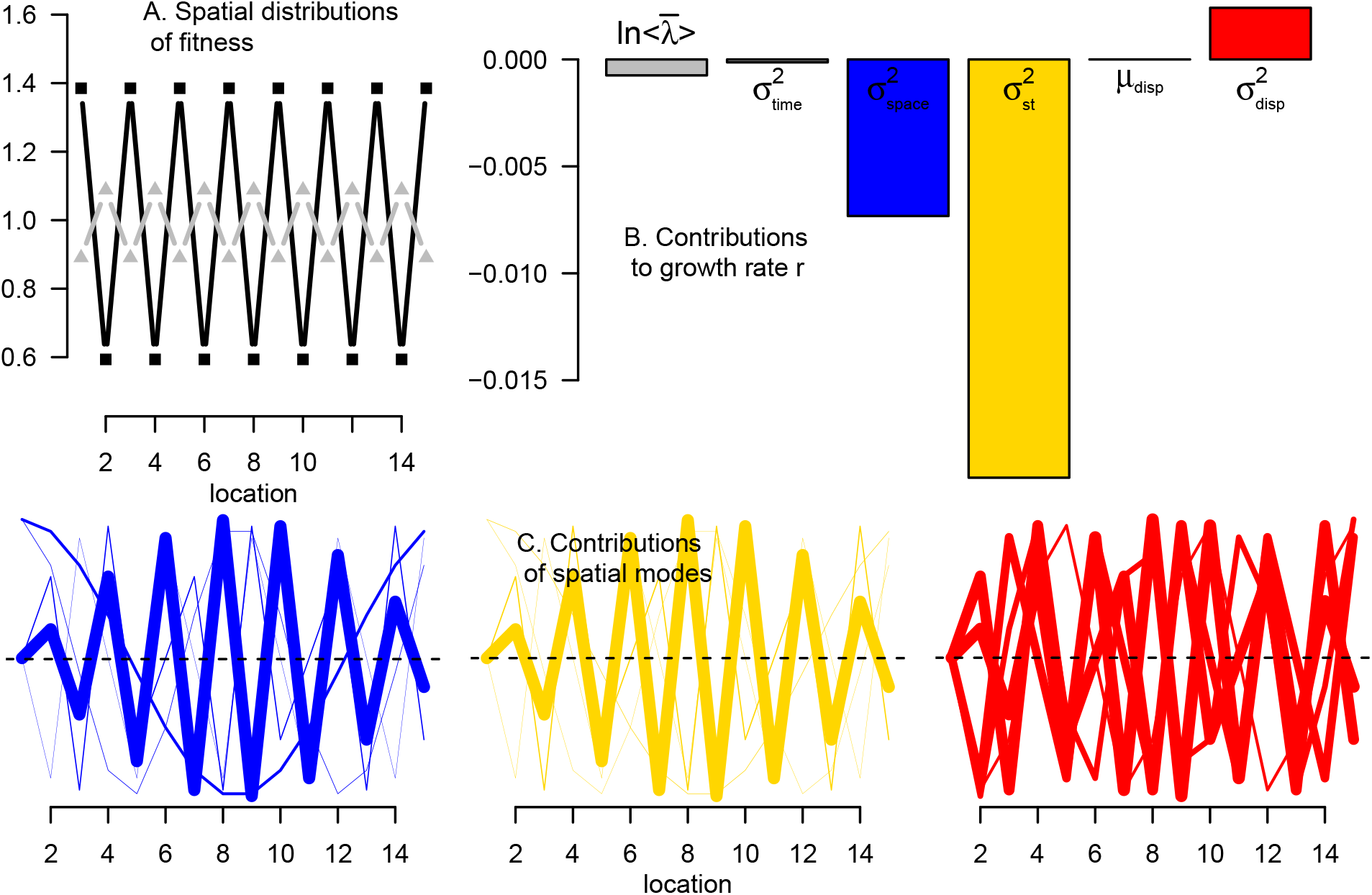
The impact of high frequency spatial variation in fitness on the metapopulation growth rate. A population living along a spatial gradient of *k* = 15 patches that varies over time between higher fitness occurring at odd number patches (black curve in A) and even numbered patches (gray curve in A). Individuals disperse to neighboring patches with a preference for patches supporting higher fitness. Analytical decomposition of the population growth rate in B. In C, all spatial modes of variation (SMV) plotted with the thickness of the curves proportional to the magnitude of their contribution to the metapopulation growth rate.

When spatial variation always occurs at lower frequencies (Fig. 4), this variation increases *r*; *r* = 0.0034 *>* 0 from both the analytical formula and numerical approximation despite ln 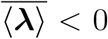 (sum of colored bars versus gray bar in Fig. 4B). While temporal variation reduces the metapopulation growth rate slightly, spatial and spatial-temporal variation in demography and dispersal more than compensate (colored bars in Fig. 4B). These positive effects stem from the spatial modes corresponding to the low frequencies (thickest curves in Fig. 4C).

When spatial variation occurs at higher frequencies (Fig. 5A), it reduces the metapopulation growth rate (*r* = *−*0.025 from analytical formula, *r* = *−*0.018 from numerical approximation) (Fig. 5B). This negative impact stems from the dominant spatial modes of variation having higher frequencies (thick blue and yellow curves in Fig. 5C) and, consequently, being miseopatric. Hence, spatial and spatial-temporal variation in demography have negative impacts on metapopulation growth rates (orange and yellow bars in Fig. 5B). These negative impacts exceed the slightly positive impacts of spatial-temporal covariation in dispersal and demography (red bar in Fig. 5B).

### Relationship to the fitness-density covariance

As shown by equation (9), the metapopulation growth rate *r* is determined by the temporal variation of the spatial averages ⟨***λ***_*t*_⟩ and the temporal variation in the fitness-density covariances 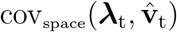. When spatial and temporal variation is small, the second-order Taylor approximation of 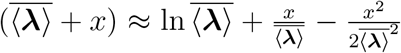 implies that the metapopulation growth rate satisfies

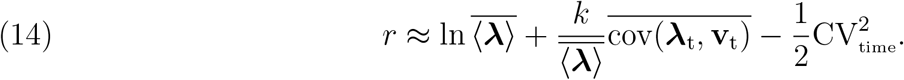

Hence, all but the first two terms of the metapopulation growth rate decomposition (13) correspond to the time-averaged fitness-density covariance. Positive values of the remaining terms (due to spatial or spatial-temporal variation in fitness, or spatial or spatial-temporal covariation in fitness and dispersal) contribute to a positive fitness-density covariance.

## DISCUSSION

Spatial and temporal variations in biotic and abiotic conditions are a ubiquitous feature of the natural world. This variation, via its effects on fitness and dispersal, modulates how quickly metapopulations grow or decline in abundance. Here, I presented a framework to explicitly quantify the contributions of each of these sources of variation to the metapopulation growth rate.

Aside from temporal variation in the spatially averaged fitness, all other sources of variation determine how population densities covary with patch quality – the fitness-density covariance of Chesson [2000]. When this covariance is positive, individuals tend concentrate in patches supporting higher fitness resulting in a higher metapopulation growth rate. This framework recovers classical results about the opposing effects of spatial and temporal variation on population growth, spatial bet-hedging by reducing temporal variation, and inflationary effects due to temporally auto-correlated variation. Furthermore, this framework highlights new mechanisms yielding inflationary effects and positive fitness-density covariances, identifies which sources of variation have the largest impact on population growth, and provides new opportunities and challenges for future theoretical and empirical work.

### Temporal versus spatial variation

A well-known dichotomy in demography is that temporal variation in fitness reduces population growth rates [Lewontin and Cohen, 1969, Boyce et al., 2006], while spatial variation can increase metapopulation growth rates [Karlin, 1976, Hastings, 1983, Snyder and Chesson, 2003, Schreiber and Lloyd-Smith, 2009, Crone, 2016]. For spatially and temporally variable environments, this dichotomy remains if temporal variation corresponds to the temporal variation in the spatially averaged fitnesses, and spatial variation corresponds to the spatial variation of the temporally averaged fitnesses. The decomposition of the metapopulation growth rate in equation (13) implies that temporal variation in spatial averages always reduces the metapopulation growth rate, and identifies when spatial variation in temporal averages increases the metapopulation growth rate. The positive effect of spatial variation only occurs if the population is philopatric, i.e. no more than 50% of individuals leave a patch in a time step; otherwise spatial variation has a negative impact. For these philopatric populations, the positive effect stems from individuals accumulating more quickly in the patches with higher fitness resulting in a positive fitness-density covariance [Chesson, 2000, Snyder, 2007]. For miseopatric populations where at least half of the individuals are leaving some patches, spatial variation reduces metapopulation growth rates. The simplest illustration of this negative impact occurs in a population dispersing between two patches with unequal fitnesses *λ*_1_ ≠ *λ*_2_. If individuals mostly disperse to the other patch each time step, then the population increases by a factor of *λ*_1_*λ*_2_ over two time steps. Hence, the metapopulation growth rate equals the logarithm of the geometric mean 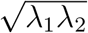. As the geometric mean is strictly smaller than the spatially averaged fitness 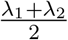, spatial variation in fitness reduces this miseopatric population’s growth rate. More generally, the metapopulation growth rate decomposition in equation (13) implies that the positive impact of spatial variation decreases continually with increasing dispersal rates i.e. the 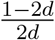 term in (10). For models without temporal variation, this prediction follows from Karlin’s reduction principle [Karlin, 1976, Hastings, 1983, Kirkland et al., 2006, Altenberg, 2012a].

The decomposition of the metapopulation growth rate clarifies under what conditions spatial variation has a larger impact on metapopulation growth rates than an equivalent amount of temporal variation i.e. *σ*_time_ = *σ*_space_. This occurs whenever dispersal rates are sufficiently low (e.g. 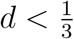 for two patch systems), otherwise temporal variation has a larger impact. This theoretical prediction is consistent with a demographic analysis of the relatively sedentary, perennial prairie forb *Pulsatilla patens* [Crone, 2016]. Using data from a 10-year demographic monitoring study, Crone [2016] analyzed two versions of an integral projection model: a spatially explicit model accounting for between-site variation in the vital rates and a stochastic model accounting for between-year variation in the vital rates. Despite spatial variation in vital rates being smaller than temporal variation in the vital rates, Crone [2016] found spatial variation had a larger effect (+0.006) on the population growth rate than temporal variation (*−*0.002).

### Spatial buffering and inflationary effects

Spatial-temporal variation, here, corresponds to the residual variation in fitness beyond spatial variation and temporal variation. This definition of spatial-variation coincides with recent work by Johnson and Hastings [2023] and Kortessis et al. [2023]. Spatial-temporal variation always corresponds to complete spatial asynchrony in fitness i.e. all spatial modes of variation have entries of opposite sign. By reducing the amount of temporal variation (in the spatially averaged fitnesses), this spatial asynchrony buffers population growth against the adverse effects of temporal variation – a classical result in metapopulation theory [Hanski, 1999]. This buffering effect is most easily understood for a two patch system with each patch exhibiting the same distribution of fitnesses over time. If the two patches are perfectly spatially synchronized, then fitness only varies over time, not space; there is a reduction of the metapopulation growth rate. If the two patches are always completely, spatially asynchronous, then fitness only varies over space, not time. When the spatial-temporal variation is temporally uncorrelated, this spatial asynchrony buffers the metapopulation growth rate by shifting all of the temporal variation to spatial-temporal variation. This well-known buffering effect can select for dispersal [Kuno, 1981, Metz et al., 1983, Levin et al., 1984], lead to evolutionarily stable sink populations [Holt, 1997, Jansen and Yoshimura, 1998, Schreiber, 2012], and rescue metapopulations from extinction [Jansen and Yoshimura, 1998, Schreiber, 2012, Evans et al., 2013]. This buffering effect, however, does not generate a fitness-density covariance.

Beyond its buffering effect, spatial-temporal variation in fitness can inflate metapopulation growth rates. Consistent with earlier work [Roy et al., 2005, Schreiber, 2010], positive autocorrelations in spatial-temporal variation increase the metapopulation growth rates of philopatric populations. Positive autocorrelation, here, corresponds to one configuration of spatial asynchrony (e.g. fitness in patch 1 being greater than fitness in patch 2) being maintained for several time steps before shifting to a different configuration of asynchrony (e.g. fitness in patch 2 being greater than fitness in patch 1). This positive effect however, is always less than the positive effect due to a comparable amount of spatial variation (e.g. 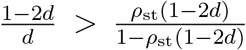 for *d <* ½ and *ρ*_st_ *>* 0), vanishes in well-mixed populations [Schreiber, 2010] and becomes negative when populations are miseopatric.

An inflationary effect also occurs for miseopatric populations when spatial-temporal variation is negatively autocorrelated. Intuitively, if patches supporting high fitness tend to have low fitness in the next time step, moving to other patches results in the metapopulation continuing to concentrate in patches with higher fitness – a positive fitness-density covariance [Chesson, 2000]. This inflationary effect likely occurs in migratory bird species dispersing seasonally between breeding grounds and wintering areas [Dingle and Drake, 2007]. Similar to the scenario considered in Figure 2, fitnesses in breeding grounds (patch 1) often vary between higher values in the summer and lower values in the winter [Lundberg, 1988, Shaw and Levin, 2011, Dingle, 2014]. In contrast, fitness in wintering areas (patch 2) are higher than the breeding grounds in the winter, but lower in the summer. If a time step in the metapopulation model is a season, then the seasonal fluctuations in fitnesses generates a negative spatial-temporal autocorrelation.

If most individuals disperse each season (*d ≈*1), then seasonal migration generates an inflationary effect. In Holarctic regions with less severe winter conditions [Lundberg, 1988], the breeding ground may support higher fitness, on average, than the wintering areas. This spatial variation can result in populations of sedentary individuals (*d ≈* 0) and populations of migratory individuals (*d ≈* 1) having locally optimal metapopulation growth rates, but populations of individuals partially dispersing (0 *< d <* 1) having low metapopulation growth rates (Fig. 2A). Hence, this combination of spatial variation and negatively auto-correlated spatial-temporal variation may contribute to the evolution of partial migration [Lundberg, 1988, Taylor and Norris, 2007, Shaw and Levin, 2011, Lundberg, 2013, De Leenheer et al., 2017] where one subpopulation always remains in the breeding grounds (*d* = 0) and the rest of the population migrates (*d* = 1).

The metapopulation growth decomposition in equation (13) also highlights how spatialtemporal variation in dispersal generates inflationary effects. This occurs when spatial-temporal variation in dispersal positively covaries with spatial-temporal variation in fitness i.e. the final term in (13). Intuitively, a tendency of individuals to track patches that will support higher fitness increases the metapopulation growth rate by creating a positive fitness-density covariance. Northern pike (*Esox lucius*) in Lake Windermere exhibit this type of positive covariance [Haugen et al., 2006]. Over a 40 year period, year-to-year movement between the southern and northern basins were positively correlated to the fitnesses in the two basins. One behavioral strategy that produces this positive fitness-dispersal covariance is the win-stay, lose-shift (WSLS) strategy. Individuals playing this strategy only leave patches after experiencing low fitness. The metapopulation growth rate decomposition (13) implies that this strategy generates a positive covariance only when the spatial-temporal variation in fitness is positively autocorrelated, a finding consistent with earlier simulation studies [Schmidt, 2004]. However, in negatively autocorrelated environments, the WSLS can enhance metapopulation growth rates provided there is pure spatial variation in fitness. WSLS can lead to individuals, on average, spending more time in these higher quality patches and, thereby, still produce a positive fitness-density covariance.

### Challenges and opportunities: Theoretical and empirical

The decomposition of the metapopulation growth rate offers many challenges and opportunities for future theoretical and empirical work. From the theoretical side, I considered environmental stochasticity, but not density-dependence or demographic stochasticity. For models with density-dependence, the partitioning of the metapopulation growth rate at low density (i.e. at the extinction equilibrium) determines which sources of variation are most important for metapopulation persistence [Benaïm and Schreiber, 2009, Benaïm and Schreiber, 2019]. However, when the metapopulation persists, it is unclear under what conditions, if any, the partitioning of the low density growth rate is predictive of the long-term spatial and temporal distribution of the metapopulation densities. Due to subtle interplay between environmental fluctuations and population dynamics, this predictive ability likely is greatest when density-dependence is compensatory but not over-compensatory [Petchey et al., 1997, Schwager et al., 2006].

In the face of demographic stochasticity and negative density-dependence, extinction in finite time is inevitable [Schreiber, 2017]. Hence, the question is not whether the population goes extinct, but how soon does it go extinct? Recently, Prodhomme and Strickler [2021] showed that a positive metapopulation growth rate *r* ensures that the time to extinction increases like a power of the total habitat size *A* i.e. *A*^*θ*^ for some *θ >* 0. In contrast, a negative metapopulation growth rate *r* implies that the time to extinction only increases logarithmically with the habitat size i.e. log *A*. Hence, in large habitats, a positive metapopulation growth rate *r* implies long persistence times, but not otherwise. Despite this fundamental dichotomy, more positive metapopulation growth rates need not, in general, imply longer persistence times or higher establishment probabilities [Schreiber and Lloyd-Smith, 2009, Pande et al., 2020a, Ellner et al., 2020, Pande et al., 2020b]. This issue stems from the metapopulation growth rate *r* only describing the mean rate of increase and not variation around this mean trend. For non-spatial models, Pande et al. [2022] introduced new metrics that account for this variation and do a much better job of predicting establishment success. To what extent these or related metrics can be used to identify the relative contributions of spatial and temporal variation to persistence times remains to be seen.

The results presented here offer many empirically testable hypotheses. Three of these hypotheses concern two-patch systems for which patch-specific fitnesses and dispersal rates can be manipulated e.g. the paramecium experiments of Matthews and Gonzalez [2007]. When one patch consistently has consistently higher fitness than the other patch, our theory predicts that the metapopulation growth rate at low dispersal (i.e. *d <* 0.5) will be higher than the growth rate for a well-mixed population, but the growth rate at high dispersal (*d >* 0.5) will be lower than the metapopulation growth rate for a well-mixed population. For the well-mixed population, one could either manipulate dispersal to achieve individuals experiencing average fitness conditions (i.e. *d* = 0.5) or manipulate the fitness in the patches to correspond to the average fitness of the spatially variable treatments. The second hypothesis concerns negative autocorrelation in space and time; one patch alternates between high and low fitness, while the other patch alternates between low and high fitness. Theory predicts that metapopulation growth rate at low dispersal rates will be lower than the growth rate for a well-mixed population, but the metapopulation growth rate at high dispersal rates will be higher than the growth rate for a well-mixed population; this corresponds to the direct opposite of the first hypothesis. The third hypothesis concerns combining the features of the first two hypothesis; the system exhibits spatial variation and negatively-autocorrelated spatial-temporal variation. There are several ways to combine these features. For example, one patch could have, on average, the higher fitness (*λ*_1_ *> λ*_2_) and the patches could exhibit negatively autocorrelated fluctuations around these averages 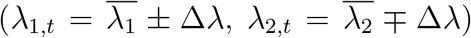. Alternatively, one could create a variable source patch and constant sink patch as described in Figure 2. Theory predicts that the metapopulation growth rates for low and high dispersal will be higher than the growth rate of a well-mixed population; this hypothesis correspond to one part of each of the first two hypotheses. Beyond these scenarios with constant dispersal, it would be exciting to test these hypothesis for different forms of variable dispersal e.g. win-stay, lose-shift dispersal versus lose-stay,win-shift dispersal.

The decomposition of the metapopulation growth rate is conceptually similar (but simpler) to the low-density growth rate decompositions in Chesson’s quantitative coexistence theory [Chesson, 1994, 2000, 2018]. Applications of Chesson’s theory to empirical systems have allowed ecologists to answer “why do these species coexist?” by evaluating the relative importance of alternative coexistence mechanisms [Adler et al., 2010, Ellner et al., 2016, 2018, Spaak and Schreiber, 2023]. Similarly, applying the metapopulation growth rate decomposition (13) to empirical systems, one could answer “why is this metapopulation growing (or declining)?” by evaluating the relative importance of different sources of variation in demography and dispersal. For example, is a metapopulation growing mostly due to uniformly high quality patches or individuals actively moving to higher quality patches? Is a metapopulation declining primarily due to temporal variation in fitness or maladaptive movement to lower quality habitats?

To answer these questions for a specific system, one needs to parameterize a spatially explicit model similar to (2). If the model involves only two patches, the symmetric dispersal matrix *D*_sym_ is given by (6) with 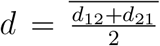. If the model involves more than two patches, there is no canonical choice of *D*_sym_, but a natural option, when possible, is using 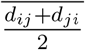 for the off-diagonal elements. Using the model, there are two approaches to decomposing the metapopulation growth rate. The computationally less expensive approach is akin to using of the second order approximation formulas of Tuljapurkar [1990] to estimate the population growth rate [Morris and Doak, 2002, Crone et al., 2013, Crone, 2016]. In our setting, one would simulate (or directly use empirical) time series of the fitnesses *λ*_*i,t*_ and the dispersal rates *d*_*ij,t*_, decompose these time series with respect to the spatial modes of variation (as described in the methods), compute the appropriate variances and covariances, and directly evaluate the decomposition of the metapopulation growth rate (13). While there is precedent for this approach in stochastic demography [Morris and Doak, 2002, Boyce et al., 2006, Crone et al., 2013, Crone, 2016], it may produce spurious results if some sources of variation are not sufficiently small.

To deal with larger sources variation in dispersal and demography, one can follow the approach of Ellner et al. [2020] for Chesson’s quantitative coexistence theory. This involves computationally more intensive calculations using a multi-step, hierarchical procedure and is based on the fitness-density covariance representation of *r* in equation (9). First, one partitions the metapopulation growth rate *r* into three terms: growth ln 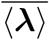 due to average fitness, a reduction Δ*r*_time_ due to temporal fluctuations, and the remaining growth Δ*r*_FD_ due to fitness-density covariance:

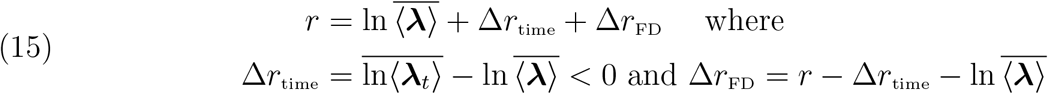

where *r* is approximated numerically as described in Models and Methods. The next step would partition the growth Δ*r*_FD_ due to the fitness-density covariance into contributions due to spatial and spatial-temporal variation in demography and dispersal. For example, if there is only variation in demography, Δ*r*_FD_ decomposes into growth Δ*r*_FD,space_ due to spatial variation and growth Δ*r*_FD,space-time_ due to spatial-temporal variation:

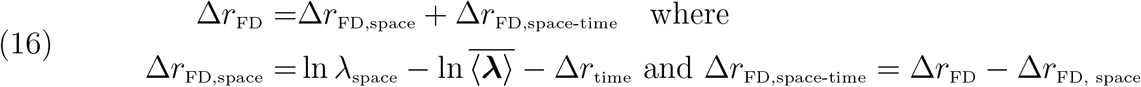

where *λ*_space_ is the dominant eigenvalue of the temporally averaged matrix 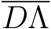.

Applying one or both of these approaches to empirically based models could provide insights into what sources of variation have the greatest impacts on the growth rates of real populations. Moreover, identifying the most important sources of variation for population growth may be useful for conservation and management efforts.

## APPENDIX A. General Formulas for First and Second Order Derivatives OF Lyapunov Exponents

This appendix derives formulas for first order and second derivatives of the top Lyapunov exponent of a random product of a stationary sequence of positive matrices. I follow the approach of Ruelle [1979] who proved under what conditions these derivatives exist and how to derive them directly using multivariate calculus. As Ruelle’s approach is presented in a skew-product formulation that can obfuscate the heart of the calculations, I use more standard probabilistic notation. Moreover, the derivations below correct an error in Ruelle’s formula for the second order derivatives, and provide explicit formulas for mixed second order derivatives not found in the classical stochastic demography literature [Tuljapurkar, 1990].

To describe the general class of random matrix products, let the set Θ of all environmental states be a compact metric space and let *θ*_1_, *θ*_2_, *θ*_3_, … be an ergodic stationary sequence taking values in Θ. *θ*_*t*_ is the state of the environment at time *t*. Let **A**(*θ*) be a non-negative, primitive matrix associated with environmental state *θ*. Assume that the mapping *θ 1→* **A**(*θ*) is continuous. **A** and *θ*_*t*_ defines a stationary sequence of matrices *A*_1_ = **A**(*θ*_1_), *A*_2_ = **A**(*θ*_2_), … and a corresponding matrix model

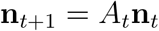

Associated with this linear model is the Lyapunov exponent

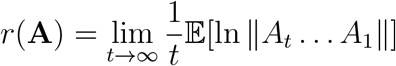

for which

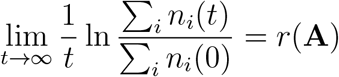

with probability one whenever the total initial population size is positive i.e. ∑_*i*_ *n*_*i*_(0) *>* 0. Furthermore, for any choice of positive row and column vectors, **v** and **w**, Ruelle [1979, Proposition 3.2] implies that

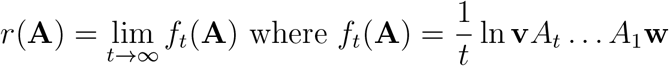

Under suitable conditions, Ruelle [1979, Theorem 3.1] proved that *r*(**A**) is an analytic function of **A**. In particular, it is continuous and continuously differentiable with respect to the environment-demography mapping **A**. Moreover, the first derivative at **A** in the **B** direction is

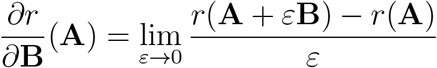

where *θ* ↦ **B**(*θ*) is a continuous mapping into the space of *k × k* matrices. Ruelle proved that

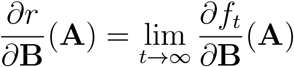

The second order derivative in the **B** and **C** directions is

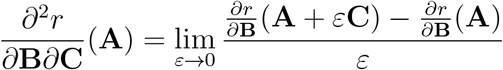

where *θ ↦* **B**(*θ*), *θ ↦* **C**(*θ*) are continuous mappings into the space of *k×k* matrices. Ruelle proved that

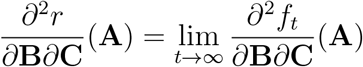

To compute 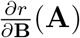, the chain rule and the product rule imply

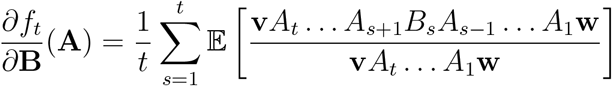

In the case of interest when **A**(*θ*) = *A* for all *θ* ∈ Θ, **v***A* = *λ***v**, *A***w** = *λ***w** and **vw** = 1, then

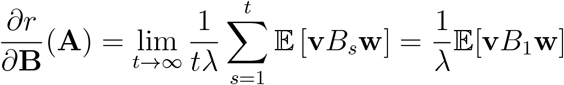

where the second equality follows from stationarity of *B*_*t*_. Thus,

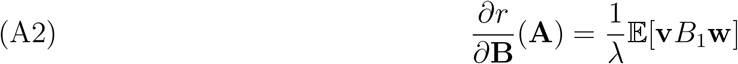

To compute a second order derivative in the directions **B** and **C**, the chain rule and product rule imply

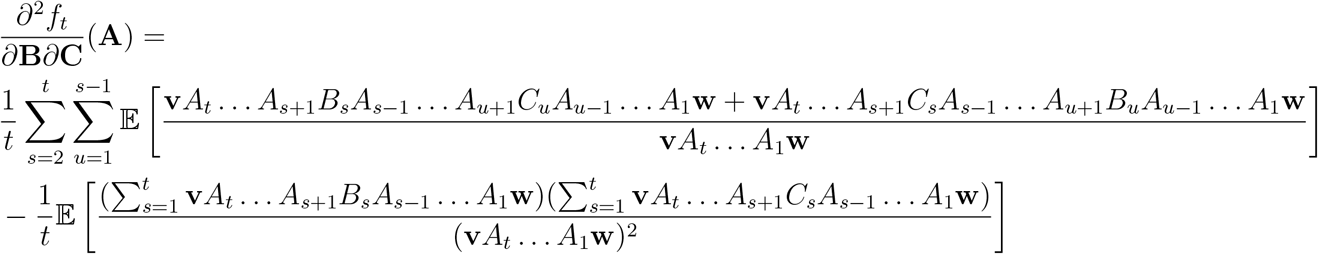

In the case of interest when **A**(*θ*) = *A* for all *θ* ∈ Θ and **v***A* = *λ***v** and *A***w** = *λ***w** and **vw** = 1, then

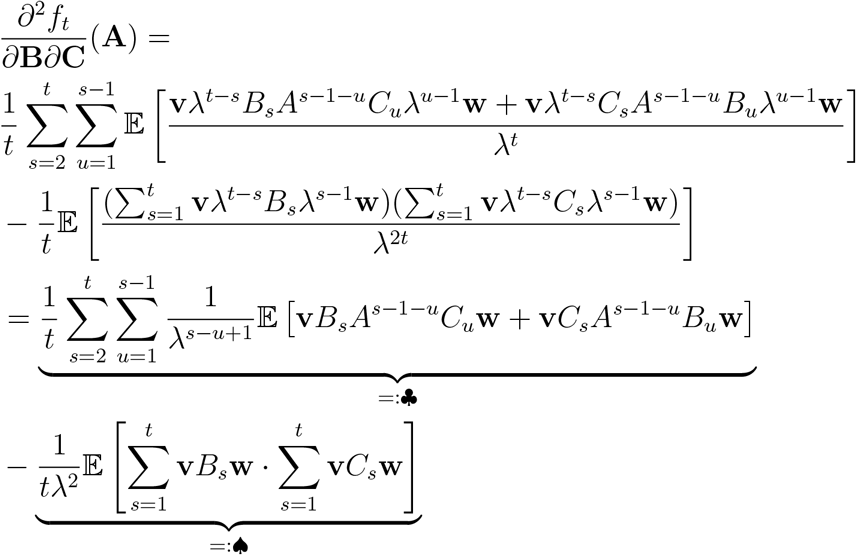

By stationarity,

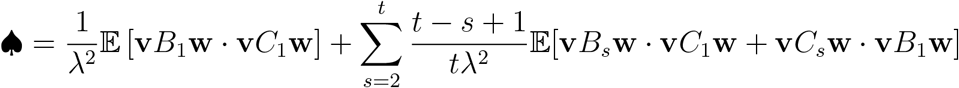

By ergodicity,

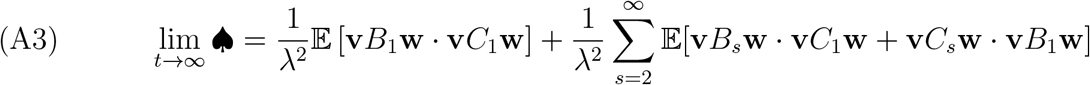

Now for ♣. Reordering the sum yields

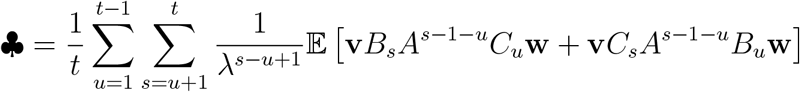

Re-indexing with *i* = *s − u* + 1 yields

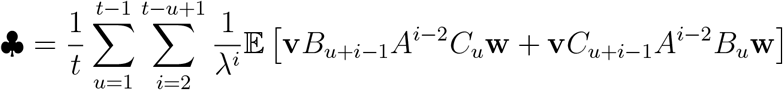

By stationarity,

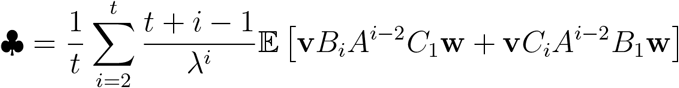

By ergodicity

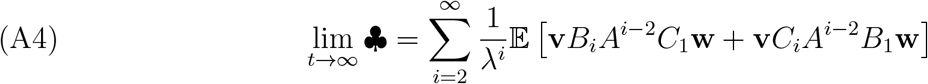

Hence, (A3) and (A4) imply (A5)

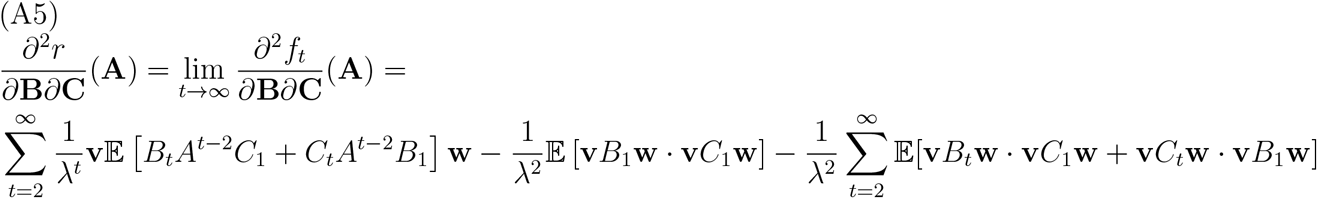

In the special case **C** = **B**, one has

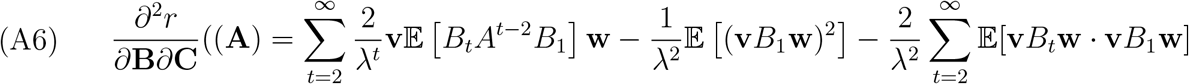

## APPENDIX B. The Metapopulation Growth Rate Approximation

In this Appendix, I derive the metapopulation growth rate decomposition (13) using the first and second order derivative formulas from Appendix A. Recall, that the metapopulation model is

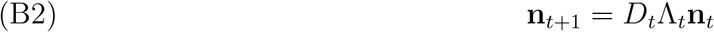

where *D*_*t*_ = **D**(*θ*_*t*_) is the dispersal matrix, Λ_*t*_ = Λ(*θ*_*t*_) is the diagonal matrix of fitnesses ***λ***_*t*_ = ***λ***(*θ*_*t*_), and *θ*_*t*_ is an ergodic, stationary sequence in the environmental state space Θ.

As described in the methods, the fitness vector can be written as 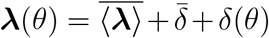 where 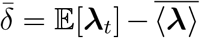 is the vector of spatial deviations, and *δ*_*t*_ = *δ*(*θ*_*t*_) is the sum of the temporal and spatial-temporal deviations. Specifically, ⟨*δ*_*t*_ ⟩ and *δ*_*t*_ *−*⟨*δ*_*t*_⟩ are the temporal and spatial-temporal deviations at time *t*. The dispersal matrix **D**(*θ*) is given by 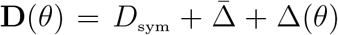 where *D*_sym_ is a symmetric, column stochastic matrix, 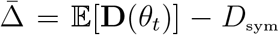 is the spatial deviation, and 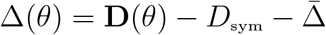 is the spatial-temporal deviation.

To use the derivative formulas in Appendix A, define the unperturbed matrix model as the constant matrix 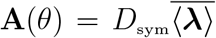 for all environmental states *θ* ∈ Θ. There are two types of perturbations to this base matrix model. First, the fitness perturbation is 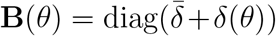. Second, the dispersal perturbation is 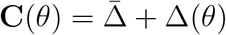 Using these perturbations, one has

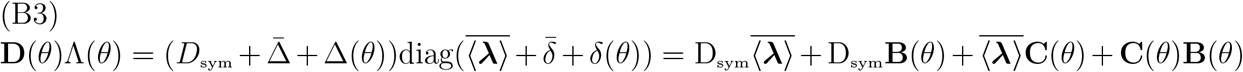

where diag(**v**) denotes a diagonal matrix whose diagonal entries are given by the vector **v**. To understand the effects of these perturbation, I compute all the relevant first and second order derivatives i.e. 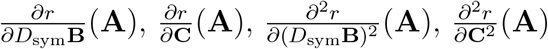 and 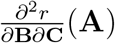.

I begin by computing the first and second order derivatives of *r* with respect to *D*_sym_**B**. Let **1** be *k ×* 1 column vector of ones, and let **1**^*T*^ denote its transpose i.e. the 1 *× k* row vector of ones.

Let **0** be the *k ×* 1 vector of zeros. Recall, that 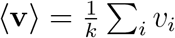 corresponds to the spatial average of a vector **v**. Using equation (A2) from Appendix A with 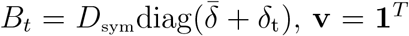 and **w** = **1***/k* yields

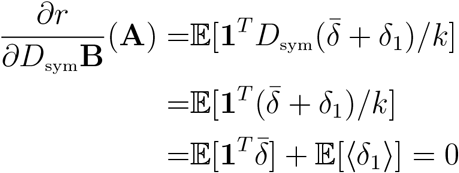

where the second line follows from **1**^*T*^ *D*_sym_ = **1**^*T*^, and the last line follows from the average spatial deviation, 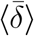, being zero, and the vector of temporal deviations having zero expectation i.e. 𝔼[*δ*_1_ ] = **0**. To find the second order derivative 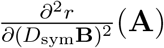, I expand the three terms in (A6) from Appendix A. For these expansions,

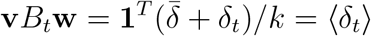

as 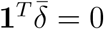. Therefore

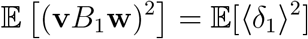

and

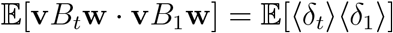

Furthermore, as 𝔼 [*δ*_*t*_] = **0**,

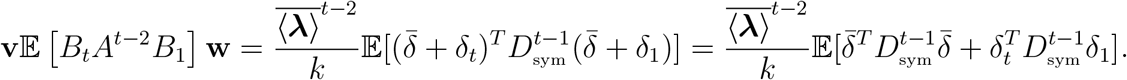

Substituting these expressions in (A6) from Appendix A yields

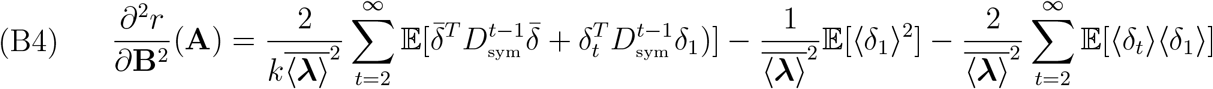

As *D*_sym_ is symmetric, one can choose a basis of eigenvectors **u**(1), …, **u**(*k*) with associated eigenvalues *μ*_1_, …, *μ*_*k*_ such that **u**(*i*)^*T*^ **u**(*i*) = *k* for all *i*, **u**(*j*)^*T*^ **u**(*i*) = 0 for *i ≠ j*, **u**(*k*) = **1** and *μ*_*k*_ = 1. **u**(1), …, **u**(*k −* 1) correspond to what the main text calls the spatial modes of variation (SMV). Let 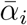 and *α*_*i,t*_ be such that

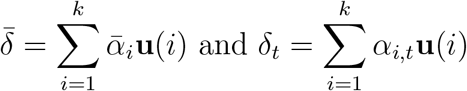

where 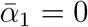. Then (B4) becomes

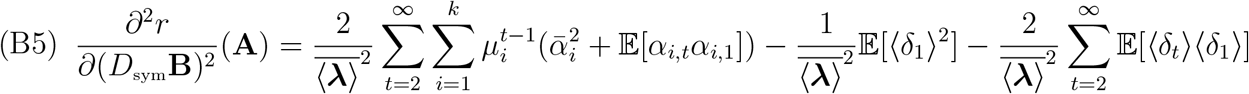

To simplify this expression further, notice that

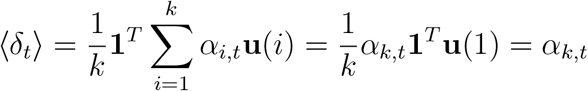

and, consequently,

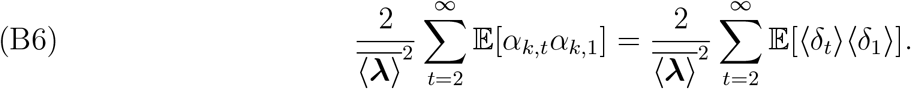

Assume 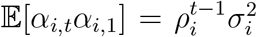 for *t ≥* 1 i.e. there is an exponential decay of correlations. Then using the fact that 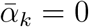 and (B6), equation (B5) becomes

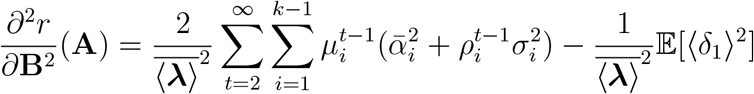

and simplifies to

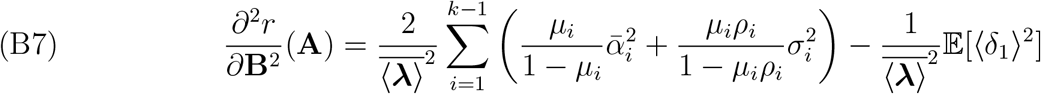

Next, I compute the derivatives of *r* with respect to **C**. As 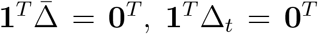, and 𝔼[Δ_*t*_] = **0**, one gets 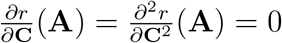. Therefore, the only contribution of the perturbation **C** is through the mixed derivative 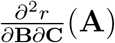. To compute this mixed derivative, I compute the terms in equation (A5) of Appendix A. To this end, notice that

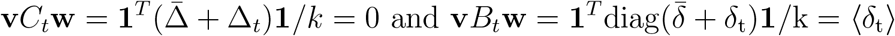

Furthermore,

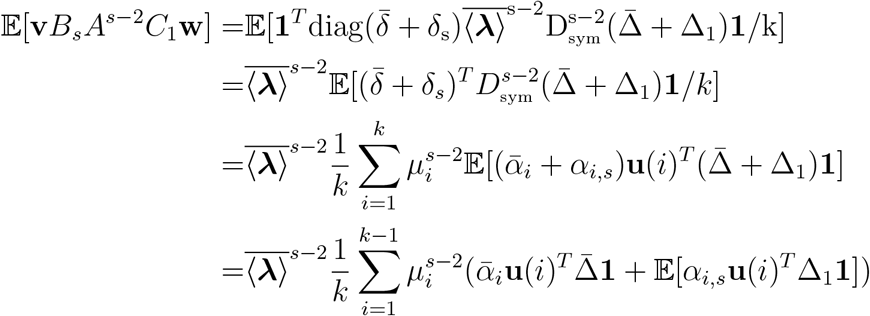

where the last equality follows from 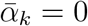, 𝔼 [*α*_*i,t*_] = 0 and **u**(*k*)^*T*^ Δ_*t*_ = 0. On the other hand,

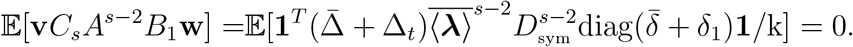

Substituting these expressions in equation (A5) yields

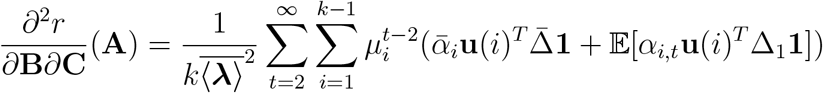

Choose 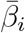 and *β*_*i,t*_ such that

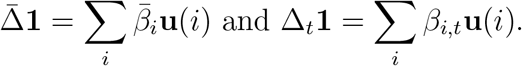

Then,

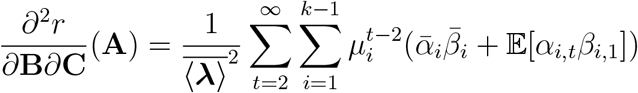

Assuming 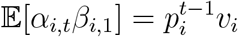 yields

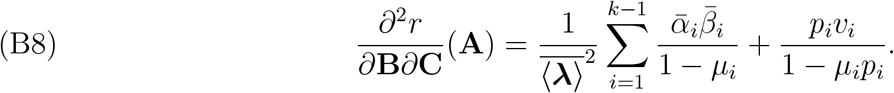

Using these derivatives and equation (B3) yields the desired second order approximation

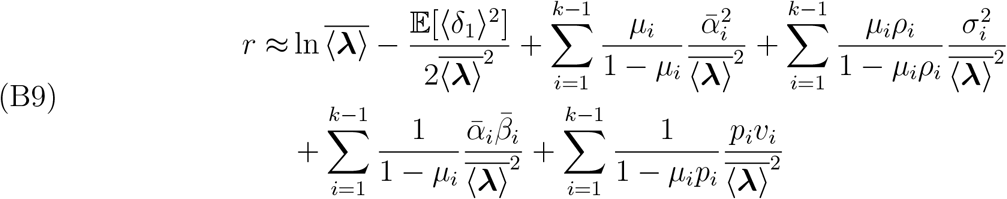

Equation (B9) gives equation (13) by defining 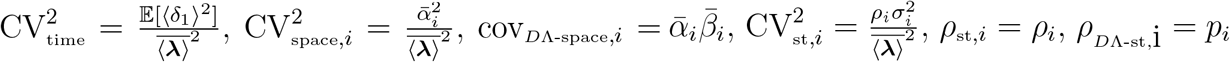, and 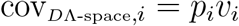.

